# Identification of Low Population States in Cryo-EM Using Deep Learning

**DOI:** 10.1101/2021.11.06.467553

**Authors:** Alec Fraser, Nikolai S. Prokhorov, John-Mark Miller, Ekaterina S. Knyazhanskaya, Petr G. Leiman

## Abstract

Cryo-EM has made extraordinary headway towards becoming a semi-automated, high-throughput structure determination technique. In the general workflow, high-to-medium population states are grouped into two- and three-dimensional classes, from which structures can be obtained with near-atomic resolution and subsequently analyzed to interpret function. However, low population states, which are also functionally important, are often discarded. Here, we describe a technique whereby low population states can be efficiently identified with minimal human effort via a deep convolutional neural network classifier. We use this deep learning classifier to describe a transient, low population state of bacteriophage A511 in the midst of infecting its bacterial host. This method can be used to further automate data collection and identify other functionally important low population states.

## Introduction

Biological structure determination by cryo-electron microscopy (cryo-EM) has become increasingly popular, comprising approximately a quarter of new submissions to the Protein Data Bank (PDB) (Berman et al., 2000). This success can be partially attributed to recent advances in sample preparation techniques, detector hardware and graphical processing unit (GPU)-accelerated software, which have altogether made structure determination increasingly attainable to experts and non-experts alike (Egelman, 2016; Kühlbrandt, 2014; Punjani et al., 2017; Zivanov et al., 2018). Secondly, the automation of cryo-EM data collection has enabled electron microscopes to collect larger datasets more efficiently, with little lag time between imaging sessions. Data collection software has now been integrated with “on-the-fly” preprocessing algorithms such as motion correction, particle picking and 2D classification (de la Rosa-Trevín et al., 2016; Punjani, 2020; Tegunov and Cramer, 2019), so that the quality of data can be characterized quickly and the microscope’s down time minimized. The future of automated data collection is “decision-free” (Li et al., 2020; Maruthi et al., 2020) cryo-EM, which will involve little-to-no human intervention and will contain high-level tools for the end-to-end analysis of cryo-EM data ranging from image acquisition to post-processing.

At present, several steps in the data processing pipeline require human intervention. Firstly, the removal of “bad” micrographs (micrograph curation) often necessitates human decision making. While it is common to simply discard micrographs based upon certain qualitative factors, such as “quality of fit” contrast transfer function (CTF) parameters, this can often lead to the inclusion of bad micrographs with good CTF parameters (Maruthi et al., 2020). Secondly, the identification of potential images of interest, or particle picking, is notoriously subjective and often requires several attempts with various software packages before the results are deemed sufficient by a human expert. Since particle picking has significant effects on all downstream data processing tasks, generating high quality particle picks is extremely important for successful cryo-EM analysis (Lawson and Chiu, 2018). To address this critical step in the cryo-EM image processing pipeline, several deep learning methods have been developed as accurate, general use particle picking procedures (Bepler et al., 2019; Wagner et al., 2019; Wang et al., 2016). However, for optimal results, these methods require a manually created training dataset prior to automated picking (Wagner et al., 2019).

Particle picking becomes even more difficult when the specimen displays preferred orientations, or the dataset contains relatively low population states of interest. In these instances, special care needs to be taken to “balance” the dataset or the resulting reconstructions will be heavily biased towards the prevalent, high population orientations and/or conformations. In the case of preferred orientations, balancing can be achieved by tilting the specimen stage and correcting for directional anisotropy (Tan et al., 2017). However, a general approach for the identification and balancing of datasets containing low-population states of interest is lacking.

In this work, we describe a method for the identification of low-population states in mixed cryo-EM datasets using a deep learning convolutional neural network (CNN). We test our method by describing the structure of a stalled sheath contraction intermediate of bacteriophage A511 that has been captured in the midst of host envelope penetration (Guerrero-Ferreira et al., 2019). We first evaluate the accuracy of the CNN classifier by making predictions on an independent validation dataset. Then, we use the classifier to identify the conformation of the particle in a larger dataset that was created by a standard template-based automated picking procedure. The accuracy of the CNN classifier is then compared to manual classification by a human expert and conventional cryo-EM classification algorithms. We conclude by describing the basis of the neural network decision making and briefly discussing the limitations and possible extensions of this method for automated data collection.

## Materials and Methods

### Bacteriophage purification

Phage A511 was propagated as described in (Guerrero-Ferreira et al., 2019). *Listeria ivanovii* strain WSLC 3009 (SV 5) was grown overnight in 1/2x Brain Heart Infusion (BHI) medium at 220 rpm at 30°C. 20 ml of an overnight bacterial culture was added to 1 L of pre-warmed (30°C) ½ BHI together with purified phage stocks to a final concentration of 10^5^ pfu/ml. This culture was incubated to an OD_600_ of ∼0.1 when additional phages were added to a final concentration of 2×10^7^ pfu/ml. Incubation continued until culture clearing for approx. 2 h when the solution was placed at 4°C. Bacterial cellular debris was removed by centrifugation at 6000×g for 15 min at 4°C. Phages were purified by the addition of 10% PEG 8.000 and 1M NaCl, overnight incubation in ice water and centrifugation at 10000×g for 15 min at 4°C. The pellet was resuspended in SM buffer (50 mM Tris-HCL pH7.5, 100 mM NaCl, 8 mM MgSO_4_) and phages were purified by CsCl gradient centrifugation (1.55 g/L) at 76000×g for 18 hours at 4°C (J et al., 2008).

### Cell wall purification

Cell wall fragments were purified and isolated from *Listeria ivanovii* as described in (Guerrero-Ferreira et al., 2019). *Listeria ivanovii* strain WSLC 3009 (SV 5) was inoculated into a 2mL ½ Brain Heart Infusion media and incubated overnight at 30°C. 1mL of the overnight culture was added to 1 L of pre-warmed (30°C) media. This culture was grown to an OD_600_ of ∼1.0 and centrifuged at 7,000xg for 10 minutes. The pellet was suspended in 10mL of 1X SM buffer (100mM NaCl, 8mM MgSO_4_, 50mM Tris-HCl pH 7.5). The sample was stored overnight at -20C. The following day the sample was thawed and heated to 100°C for 20 minutes. The cell walls were disrupted by passing the sample through a pressure cell homogenizer three times. The sample was then centrifuged at 20,000xg for 30 minutes. The pellet was suspended in 500mL water and centrifuged at 20,000xg for 30 minutes, and repeated once more. 100ug of RNAse A and DNAse 1 were added to the sample and shaken for 3.5 hours at room temperature. At this point, 100ug of Proteinase K was added and shaken for 2 hours at room temperature. The sample was boiled in 4% SDS for 30 minutes at 100°C. The sample was centrifuged at 20,000xg for 30 minutes, and the supernatant was decanted. The pellet was washed in 500mL of water for 20 minutes at 20,000xg and repeated for a total of six washes. The pellet was suspended in 10mL SM buffer and placed at 4°C for storage.

### Cryo-EM Sample preparation

Cell membranes were sonicated for 15 minutes. The bacteriophages were then incubated with the cell membranes for 3 minutes prior to grid freezing. TED PELLA 200 mesh PELCO NetMesh copper grids were plasma cleaned by the Gatan advanced plasma cleaning system. 3 ul of sample was pipetted onto the plasma cleaned grids using a Thermo Fischer Scientific Vitrobot for 20s at 100% humidity. The grids were plunged into liquid ethane.

### Data collection, image correction, and particle picking

A Titan Krios 300kV electron microscope with a BioQuantum K3 imaging filter and 20eV electron slit was used for imaging. Using EPU software, 32,260 micrographs (4096 × 4096 size, 59 frames, 1.1 Å pixel size, exposure time of 1.5s and a total electron dose of 40 e/Å^2^) were collected over a defocus range of **−**1.5 to **−**3.5 microns. The micrographs were patch motion corrected and CTF estimation was performed using cryoSPARC (Punjani et al., 2017). Micrographs whose CTF fit resolution was > 8 Å were discarded, resulting in a set of 30,069 micrographs. A 2D template, generated from 200 manually picked contracted particles, was used as an initial template in an automated particle picking procedure as implemented in cryoSPARC (Punjani et al., 2017). The resulting picks were classified using 2D classification and new templates were created for automated picking. This procedure was iterated 4 times until the quality of particle picks plateaued, resulting in 6,670 putatively contracted particles (after 2D classification).

### Creation of labelled data

A subset of the entire cryo-EM dataset was curated for manual picking and the labelling of images (**Fig. 1A**). We manually examined the first third of our dataset (∼10,000 raw images) and identified ∼2,400 fully contracted particles (**Fig. 1B**) and only 400 stalled intermediates (**Fig. 1C**). We then selected these stalled intermediates and the first 400 contracted particles for neural network training. These two 400-image sets were further separated into 318 and 82 images that comprised the training and validation sets, respectively.

**Figure 1.**
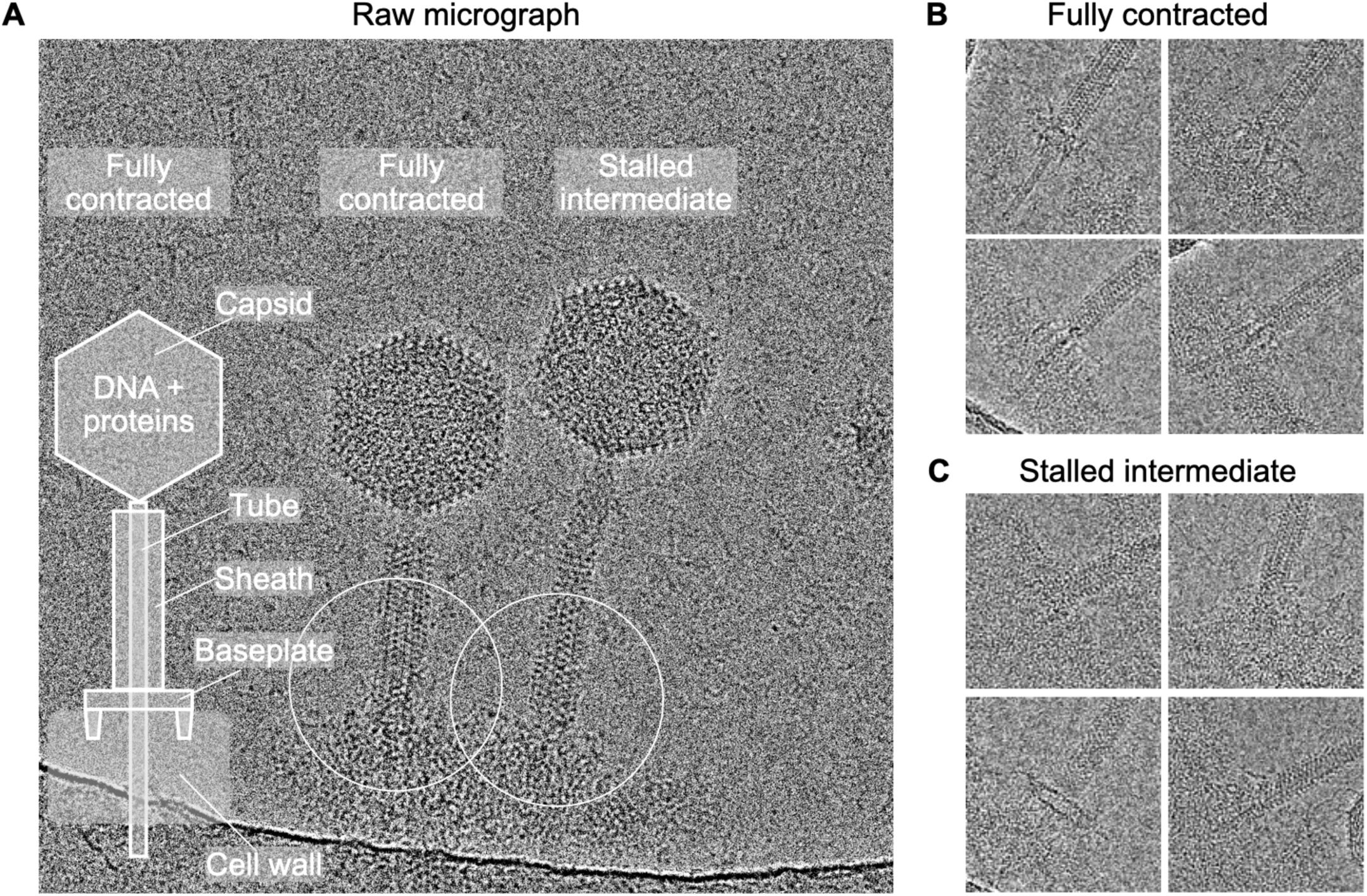
Cryo-EM Data. **A**, Raw micrograph with a diagrammatic representation of the A511 virus particle in the contracted sheath state. Raw micrograph showing both fully contracted and stalled intermediate bacteriophages. The white circles show the size of the reconstructed volume for both states. **B, C**, Representative images of the baseplate proximal sheath structure used for model training in the contracted and stalled intermediate states, respectively.

The inherently limited size of the training data is a major challenge to the implementation of deep learning techniques. For this reason, the two training sets were combined and subject to data augmentation techniques. 636 training images were rotated (using EMAN2 (Tang et al., 2007)) by 90, 180 and 270 degrees and flipped horizontally, resulting in 5,088 images. Each of these images was translated with periodic boundary conditions along both the x- and y-axes by one pixel (8 unique translations and the original non-translated image), resulting in 45,792 images.

Throughout training, model performance was evaluated by making predictions on an independent validation dataset, consisting of 164 particle images, rotated by 90, 180 and 270 degrees and flipped horizontally, resulting in 1,312 images (comprising 1/8^th^ of the manually picked dataset). Neural network input images were binned to a pixel size of 12.9 Å and a box size of 128 pixels. Images were normalized to have a mean pixel value of 0 and a standard deviation of 1. Each image had gaussian noise (mean=0, variance=0.25) added to improve the tolerance of the neural network to noise.

### Neural Network Architecture

The neural network was composed of three convolutional layers followed by two dense layers and a final softmax layer (Dunne and Campbell, 1997) (**Fig. 2**). Each of the convolutional layers was composed of 64 (15×15) filters with layer weight regularizers (van Laarhoven, 2017) (L2=5e-4). The convolutional layer was followed by a rectified linear unit (Relu) (Nair and Hinton, 2010) activation function. Batch normalization was applied to the outputs of the Relu function. 2×2 Max pooling was performed following batch normalization, shrinking the size of the images by a factor of 2.

**Figure 2.**
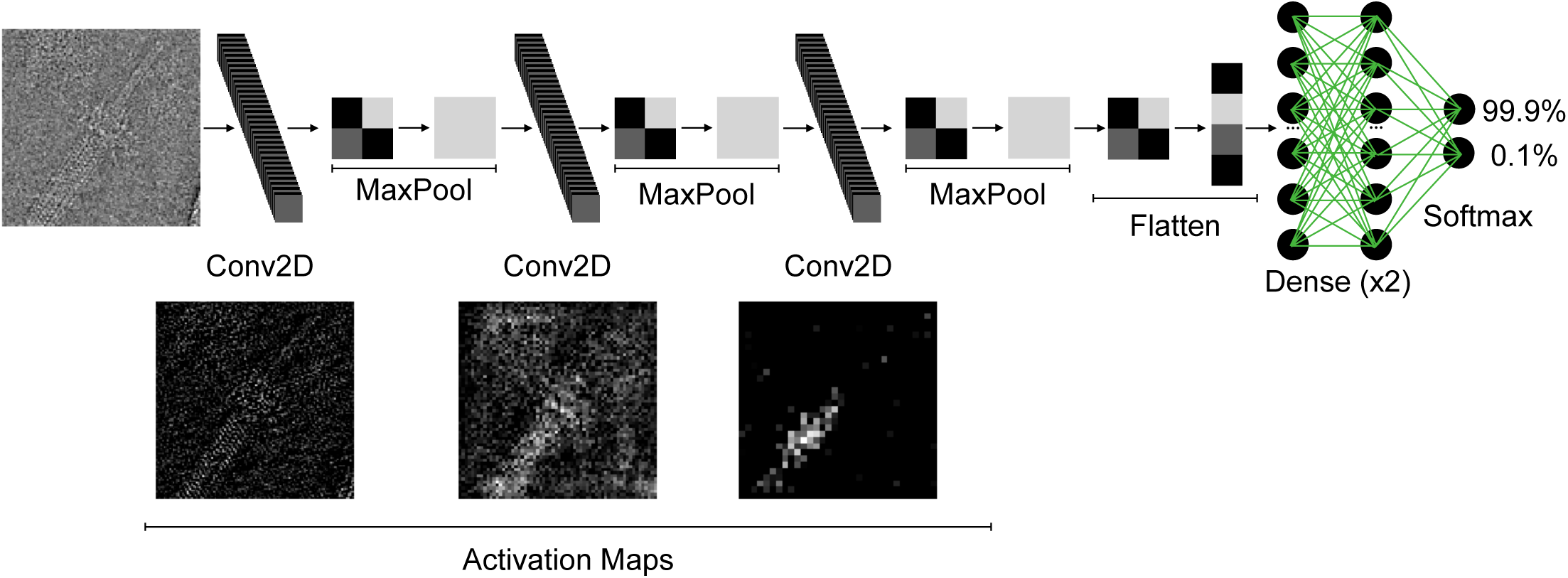
Neural Network Architecture. A diagram of the CNN in which the layers are shown simplistically for clarity. Activation, batch normalization and dropout layers are omitted for clarity. The activation maps of the first, second and third convolutional layer are shown below the CNN diagram. White pixels correspond to regions where the input to the convolutional layer is similar to the trained filter. Activation maps show image binning as a result of max pooling operations.

After the three convolutional layers, a dropout layer (rate=50%) was used to reduce overfitting. The result was then flattened prior to connection with two identical fully connected (dense) layers with 500 units. Batch normalization was applied to the outputs of the dense layer per mini-batch. Batch normalization was followed by a rectified linear unit (Relu) activation function. Finally, a dropout layer (rate=50%) was used to reduce overfitting. After these two dense layers, a final dense layer with two units and a softmax activation function was used to predict the probabilities of the two classes (stalled intermediate and contracted).

### Training

Model training was performed on a set of 45,792 128×128 particle images, which included equal numbers of contracted and stalled intermediate particles (**Fig. 3**). The model was compiled with an Adam optimizer (LR=5e-7) and a binary cross entropy loss. Training was performed for 196 epochs, at which point the validation loss and validation accuracy reached the global extremum values of 0.33 and 0.916 and training was halted **(Fig. 3A 3B**).

**Figure 3.**
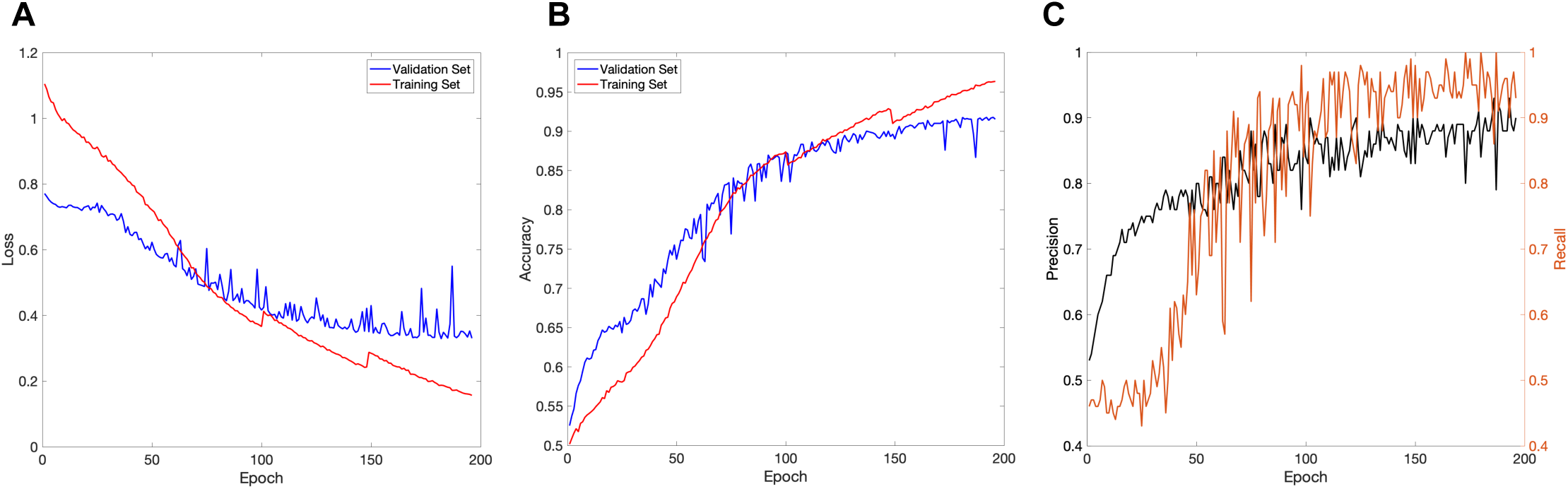
Model Training. **A**, Graph showing the model loss on the training and validation sets throughout training. **B**, Graph showing the overall model accuracy on the training and validation sets throughout training. **C**, Graph showing precision and recall on the validation set throughout training. Discontinuities in (**A, B**) around epochs 100 and 150 were caused by the restarting of model training.

### Implementation

Model training and prediction was performed using the Texas Advanced Computing Center (TACC) Stampede2 high performance computer (HPC).

### Molecular Graphics

Graphs in figures 3, 4 and 7 were created using MATLAB (MathWorks). Figure 6 was created using UCSF ChimeraX (Goddard et al., 2018). Supplementary figure 1 was created using UCSF ChimeraX (Goddard et al., 2018) and UCSF Chimera (Pettersen et al., 2004).

**Figure 4.**
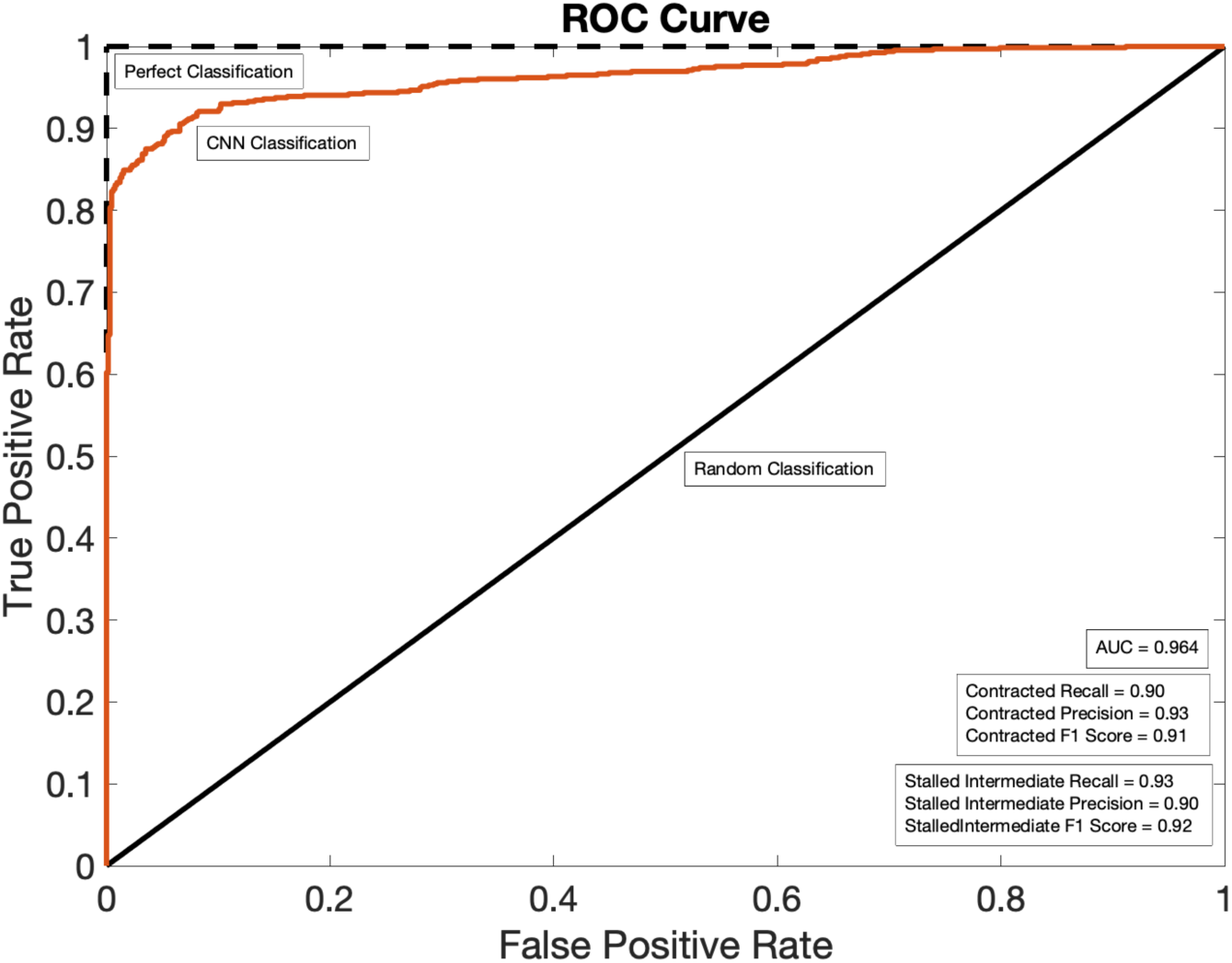
Deep Classifier Accuracy. ROC curve showing the TPR and FPR of the classifier on a validation dataset. The theoretical performances of a perfect (or ideal) and random classifier are shown as a dashed black line and a solid black line, respectively. The performance of the CNN classifier described here is shown as a red line. The recall, precision, and f1-scores for the identification of contracted and stalled intermediate images, as well as AUC, are given.

## Results

### Rationale and complexity of the problem

The host cell attachment organelle of bacteriophage A511 – its tail – undergoes a large conformational change upon host recognition and binding (Guerrero-Ferreira et al., 2019). The tail’s outer sheath contracts, propelling the rigid inner tube to pierce the host cell envelope because the base of the sheath is anchored to the envelope via a baseplate (Fraser et al., 2021). Contraction stalls in a fraction of A511 bacteriophages and, if properly timed, a mixture of fully extended (pre-attachment or just-attached), fully contracted, and stalled contraction intermediates (termed ‘stalled intermediates’) can be captured and imaged in cryo-EM (**Supplementary Fig. 1A**). While the populous fully extended and fully contracted states can be distinguished using conventional cryo-EM classification methods, low-population stalled intermediates are often wrongly classified as fully contracted or simply discarded as bad particles. As such, critical information about the propagation of the contraction wave is lost (Fraser et al., 2021). The problem of distinguishing fully contracted from stalled intermediate particles is further exacerbated by the inherent flexibility of the sheath in the stalled intermediate state (**Fig. 1A, Supplementary Fig. 1A**).

In this work, we created a deep learning CNN (**Fig. 2**) to distinguish the stalled contraction intermediate from the fully contracted sheath particle using the baseplate-proximal part of the sheath (**Fig. 1B, 1C**). Given the imbalance between the total number of fully contracted and stalled intermediate particles (∼6:1), a particle set with substantially more stalled intermediate than contracted particles is likely necessary for the successful reconstruction of a stalled intermediate electron density map, without substantially reduced resolution and map blurring (Serna, 2019). As a consequence, the CNN’s ratio of the True Positive Rate (TPR) over the False Positive Rate (FPR) must be substantially greater than six.

### Accuracy of Classification

The accuracy of the deep classifier was evaluated on an independent validation dataset which was not used for CNN training. As such, neural network performance on this dataset is representative of its performance on data which it has not seen before. The receiver operating curve (ROC) plot (**Fig. 4**) shows the relationship between the TPR and the FPR on the validation dataset. Upon integration of the ROC curve, the area under the curve (AUC) was calculated to be 0.964, close to a perfect score of 1.0, showing that the model has a good measure of separability between stalled intermediate and contracted images. The recall, precision and f1-score (Powers, 2020) for the prediction of contracted images was determined to be 0.90, 0.93 and 0.91, respectively. The recall, precision and f1-score for the prediction of stalled intermediate images was determined to be 0.93, 0.90 and 0.92, respectively.

### Generation of an Electron Density Map from CNN Classified Particles

Application of the CNN classifier to the entire dataset of 6,670 automatically picked particles resulted in the identification of 1,745 stalled intermediates (**Fig. 5**). These particles were subject to multi-class ab-initio reconstruction in cryoSPARC. The most populous and best resolved class had 995 particles and was refined with C6 symmetry to a resolution of 12.8 Å (**Fig. 6A, Fig. 7**). The resulting electron density displayed a conformation that is distinct from those of both the fully extended and contracted states, which will be described elsewhere (**Supplementary Fig. 1A, 1B, 1C**). Importantly, the quality of the map was sufficient for the unambiguous rigid docking of an atomic model derived from a 3.1 Å resolution fully contracted map of the A511 sheath (**Supplementary Fig. 1D**).

**Figure 5.**
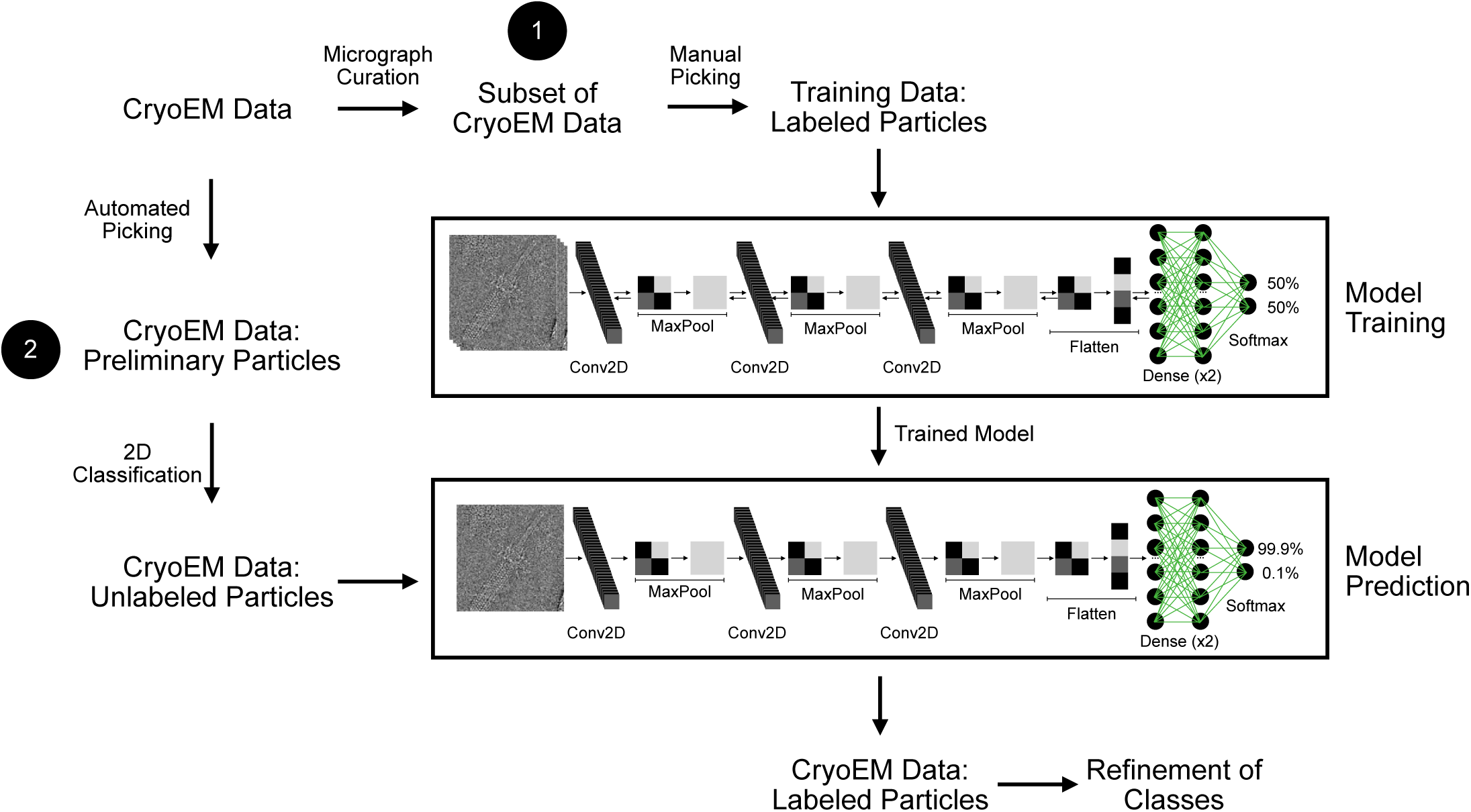
Method Workflow. Diagram showing the general procedure for the implementation of the deep learning classifier. The pathways 1 and 2 depict the training and prediction module, respectively. Forward and backward arrows in the model training image represent the updating of weights and biases throughout training. Forward arrows in the model prediction image indicate a singular pass through the network. Activation, batch normalization and dropout layers are omitted from the model training and model prediction images for clarity. Compulsory data preprocessing steps (such as motion correction and CTF estimation) are omitted from the workflow for clarity.

**Figure 6.**
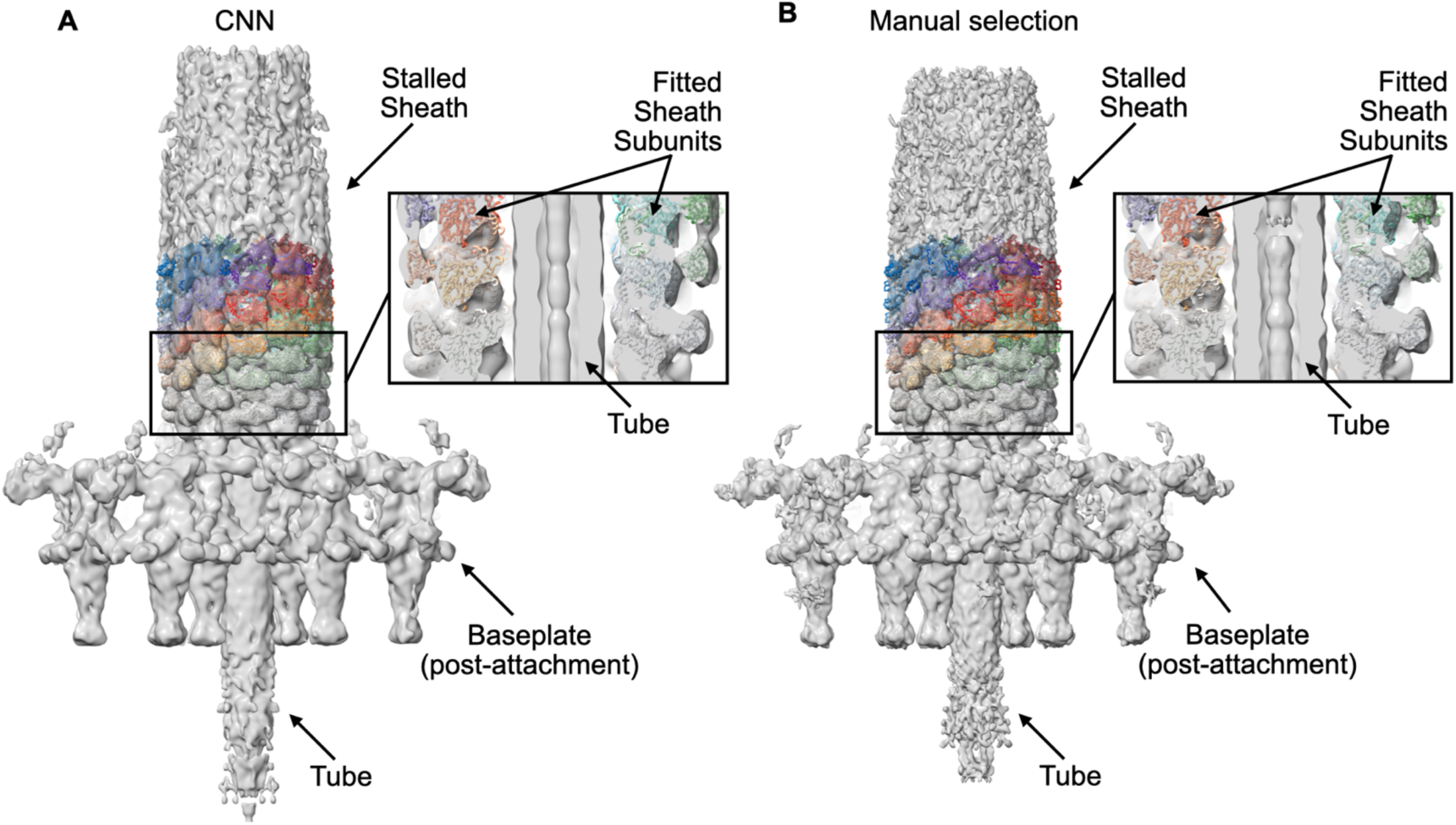
Comparison of CNN-derived and Manually Curated Electron Density Maps. **A, B**, Electron density map derived from CNN identified and manually picked particles, respectively, with fitted sheath subunits. The inset shows cut-away highlighting the fit of the sheath subunits in the electron density map. Each of the six helical strand of the fitted sheath subunits is colored in a distinct color.

**Figure 7.**
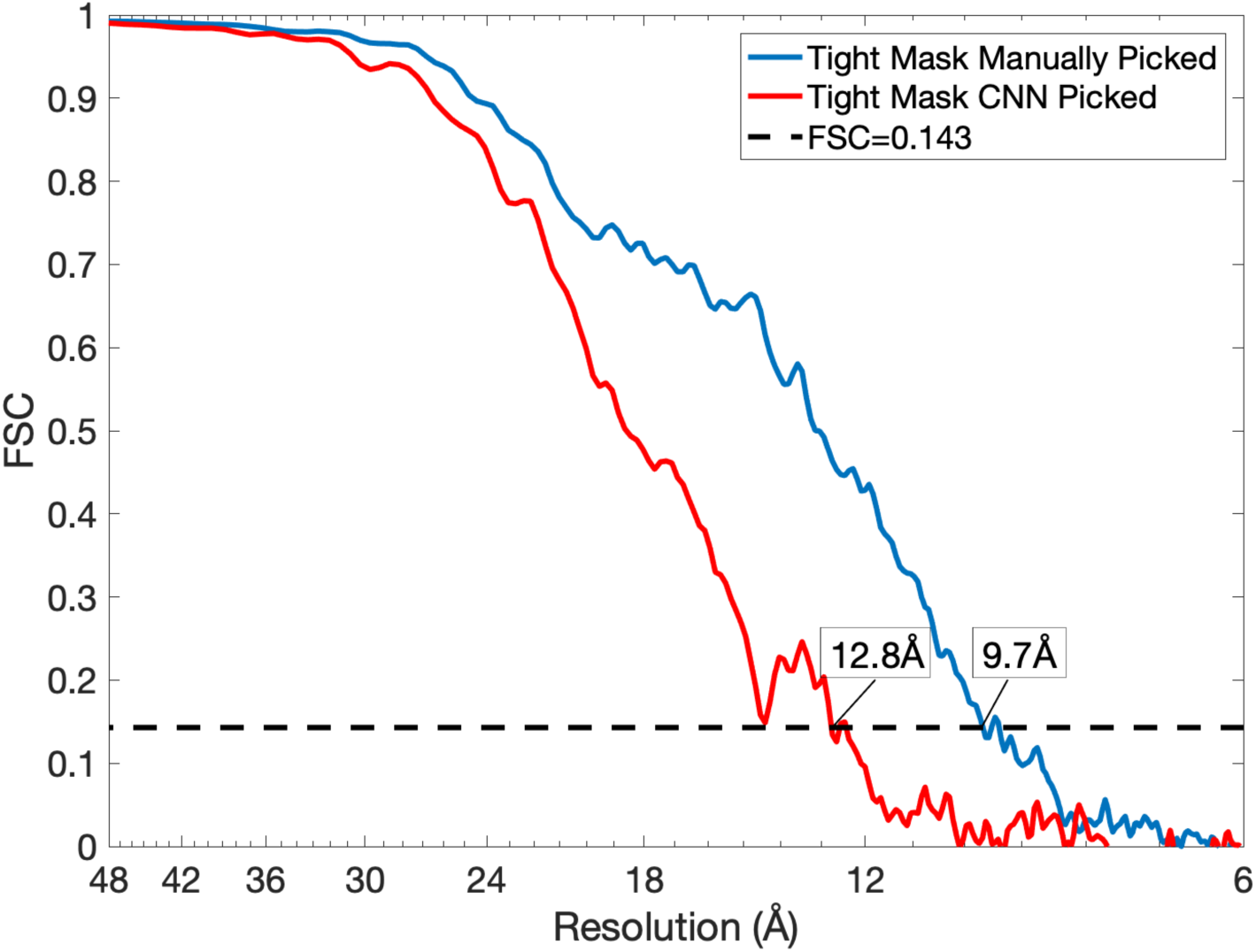
Resolution of CNN Derived and Manually Curated Cryo-EM Maps. FSC curves of the two cryo-EM maps of the stalled intermediate shown in **Fig. 6** that were calculated using manually picked particles and selected by the CNN. The dashed line corresponds to the FSC value of 0.143 and serves for the determination of nominal resolution of cryo-EM reconstructions.

### Comparison of CNN-derived to Manually Picked Dataset

Next, CNN accuracy was compared to the accuracy of a human expert. Manually curated stalled intermediate particles were extracted from the entire cryo-EM dataset, which, after 2D classification, resulted in 1,164 particles. These particles were subject to multi-class ab-initio reconstruction in cryoSPARC. The most populous and best resolved class had 814 particles and was refined with C6 symmetry to a resolution of 9.7 Å (**Fig. 6B, Fig. 7**). This electron density map had a higher resolution (**Fig. 7**) and contained more detail than the CNN-derived map (**Fig. 6**). Nevertheless, the two maps have a correlation coefficient of 0.963, which shows that both maps represent the same conformer and carry a comparable amount of functionally relevant information.

### Comparison of CNN-derived Electron Density to that Derived from Conventional Cryo-EM Classification Techniques

Conventional 3D classification techniques were used in an attempt to generate a homogenous subset of stalled intermediate particles and create a well-resolved electron density map. 6,670 automatically picked particles were used to generate various *ab initio* models. These models acted as initial models for 3D classification with 3, 5 and 7 classes. In some cases the initial models were low-pass filtered to 120 Å resolution to reduce the bias of the prevalent contracted state. In other instances, 3D classification was performed with C6 symmetry. In all other cases, a low pass filter of 20 Å was applied to the initial models and no symmetry was imposed. In all instances, the stalled intermediate state of the bacteriophage sheath remained unresolved. In some cases, blurry electron density maps, likely representing a superposition of contracted and stalled intermediate particles, were observed. However, further classification failed to produce a more homogenous subset. Ultimately, these methods were unsuccessful in generating an electron density map of sufficient quality for the unambiguous placement of sheath subunits. It is likely that a combination of low particle numbers and the predominance of contracted particles in the dataset led to the inability of 3D classification procedures to group stalled intermediate particles into a single class.

## Discussion

### Interpreting the Neural Network Decision Making

To better understand the inner workings of the CNN, we generated activation maps of validation set images (**Fig. 2**). Activation maps are a visual representation of the linear algebraic operations made on an image at a certain point in the network. The activation maps of the first convolutional layer show filters trained to detect lines and edges. The activation maps from the last convolutional layer show filters trained to detect the symmetric patterns of the bacteriophage sheath. Altogether, this is consistent with the notion that CNNs make predictions based on local features as opposed to global features (Baker et al., 2020, 2018, n.d.).

### Extension of the Method in Automated Data Collection

Future applications of this method can be used to streamline automated cryo-EM data collection. At present, automated data collection can perform routine pre-processing steps “on-the-fly”, such as motion correction, particle picking and 2D classification. In cases where particle counts are high, there exists efficient strategies whereby initial 3D reconstructions can be derived in hours (Stabrin et al., 2020). However, in cases where conformations of interest are rare and other states are more populous, established methods for automated data collection simply discard low population states, as in our case.

As an alternative, an initial set of micrographs can be used to generate training images for a deep neural network classifier. Following the generation of labeled data and subsequent model training, the CNN can be used to classify particles generated from other on-the-fly particle picking methods. After classification, micrographs which do not contain states of interest can be deleted, also on-the-fly, to reduce disk space requirements, which are especially acute when searching for low population states.

### Limitations of the Method

The ability of the deep classifier to predict stalled contraction intermediates with high accuracy using an astoundingly small training dataset is likely a consequence of the symmetric structure of the bacteriophage sheath (**Fig. 1A, Supplementary Fig. 1A, 1B, 1C**). Given the size of the sheath, each training image contains dozens of copies of either the stalled intermediate or contracted sheath pattern. As a consequence, while only 636 non-augmented images were used for training, the neural network was effectively trained on thousands of non-augmented instances of the sheath pattern. Therefore, it is possible that this method would be less effective, with such limited training data, if the object of interest possesses no symmetry. We expect that this method would work well, with similarly limited training data, for the characterization of low population states of filamentous or otherwise symmetric structures, such as the flagellum, cilium and myosin.

## Author contributions

**A.F**. and **P.G.L** conceived the study. **A.F**. processed the cryo-EM data and performed all deep learning analysis in its entirety. **J.M.M**. prepared the Listeria cell wall sample. **E.S.K** prepared the phage A511 sample. **N.S.P**., **A.F**. and **E.S.K**. optimized the sample for cryo-EM imaging. **N.S.P**. collected cryo-EM data. **A.F**. and **P.G.L** wrote the manuscript, which was read, edited, and approved by all authors.

## Competing interests

The authors declare no competing interests.

## Correspondence and requests

Correspondences should be addressed to Alec Fraser and Petr G. Leiman.

## Acknowledgements

This work was supported by the UTMB Sealy Center for Structural Biology and Molecular Biophysics, by the Department of Biochemistry and Molecular Biology, and by the National Institute of Health (Grant R01GM139034). We thank Dr. Michael B. Sherman for his help in Cryo-EM data collection. The work present in this paper used the resources of the UTMB SCSB Cryo-EM laboratory. We thank the Texas Advanced Computing Center (TACC) for their computational resources which supported this work.

## Code availability

Code for the training of the CNN can be found at: https://github.com/alecfraser94/Deep-Learning-for-the-Identification-of-Low-Population-States-in-Cryo-EM

**Supplementary Information** is available for this paper.

## Notes

https://github.com/alecfraser94/Deep-Learning-for-the-Identification-of-Low-Population-States-in-Cryo-EM

### Competing Interest Statement

The authors have declared no competing interest.

