## Supplementary Information for "Identification of Low Population States in Cryo-EM Using Deep Learning"

### Contents

**Supplementary Figure 1**

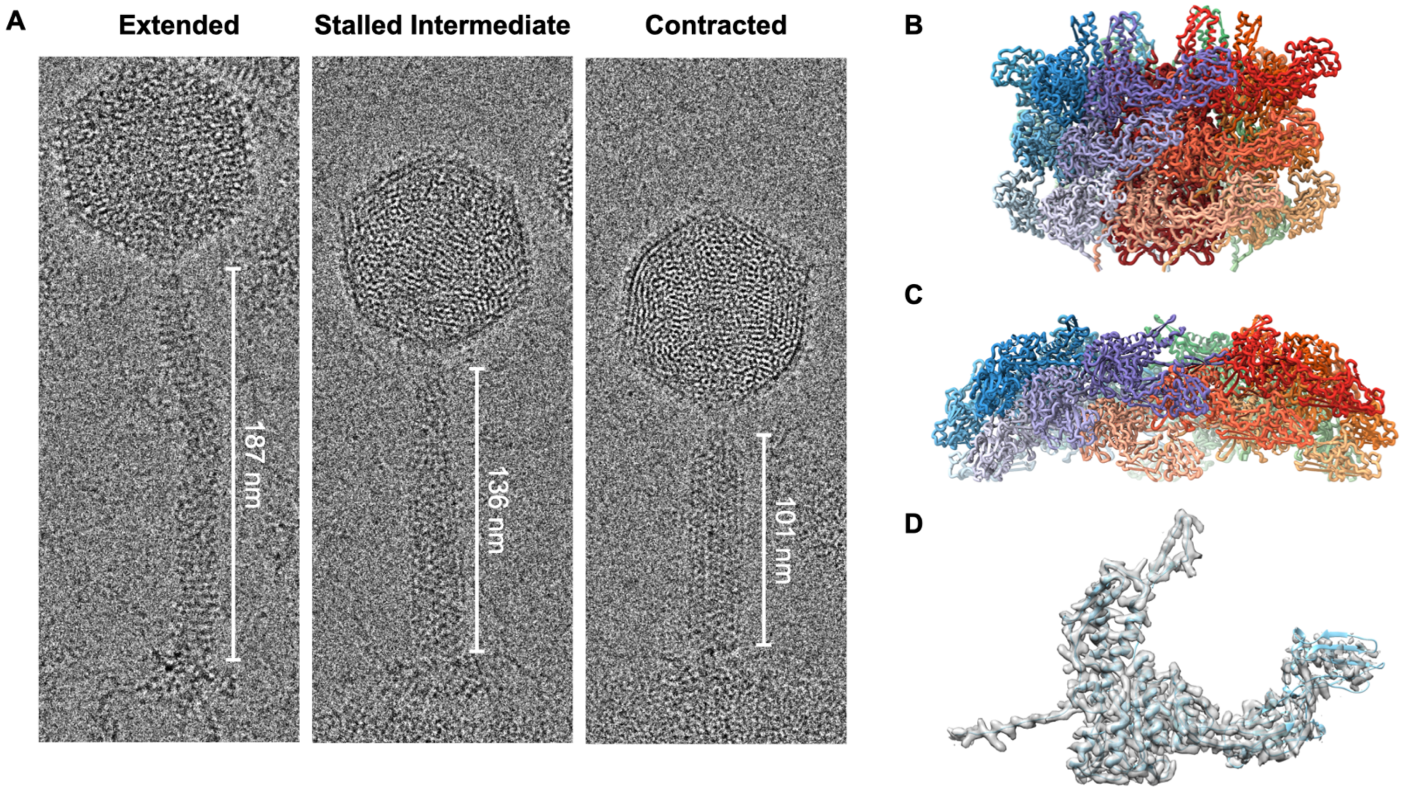


**Supplementary Figure 1. Three Conformations of the Bacteriophage A511 Tail.**

**A**, Electron micrograph images of the bacteriophage A511 in the extended, stalled intermediate, and contracted sheath states.

**B**, Ribbon diagram of three layers of the sheath-tube complex in the extended state. The sheath is colored according to its strands. The tube is colored brown.

**C,** Ribbon diagram of three layers of the contracted sheath. The sheath is colored according to its strands.

**D**, A ribbon diagram of a single subunit of the sheath in the contracted state (cyan) with its cryo-EM electron density (semi-transparent gray) at 3.1 Å resolution. This atomic structure was used in the interpretation of the stalled contraction intermediate maps shown in **Fig 6**.
